# NKCC1 a Regulator of Glioblastoma Progression

**DOI:** 10.1101/2025.09.19.677289

**Authors:** Anja Thomsen, Diana Freitag, Madlen Haase, Christian Senft, Falko Schwarz, Silke Keiner

## Abstract

Glioblastoma (GBM) is the most common and aggressive primary brain tumor in adults, with poor prognosis despite multimodal therapy. Chloride cotransporters NKCC1 and KCC2 are key regulators of intracellular chloride levels and thereby determine whether GABA acts inhibitory or excitatory. In GBM, disrupted chloride homeostasis promotes proliferation, migration, and stem-like properties, but its clinical relevance is not fully understood.

We analysed *NKCC1* and *KCC2* expression in glioblastoma samples, considering clinical parameters such as age, gender, and MGMT promoter methylation. Statistical analyses included ROC-based cutoff determination, Kaplan–Meier survival analysis and subgroup. Immunohistochemistry was performed to identify cell types expressing NKCC1.

*NKCC1* expression was significantly higher in older patients and emerged as a prognostic marker for recurrence-free survival, with lower levels correlating with delayed recurrence, although overall survival was unaffected. *NKCC1* was expressed in stem-like, astrocytic, and progenitor cells, but not in mature neurons.

These findings identify *NKCC1* as a regulator of GBM progression and recurrence, linking chloride transporter imbalance to GABAergic signalling. Targeting *NKCC1* and restoring chloride homeostasis may provide promising new treatment strategies.

**Graphical abstract:** 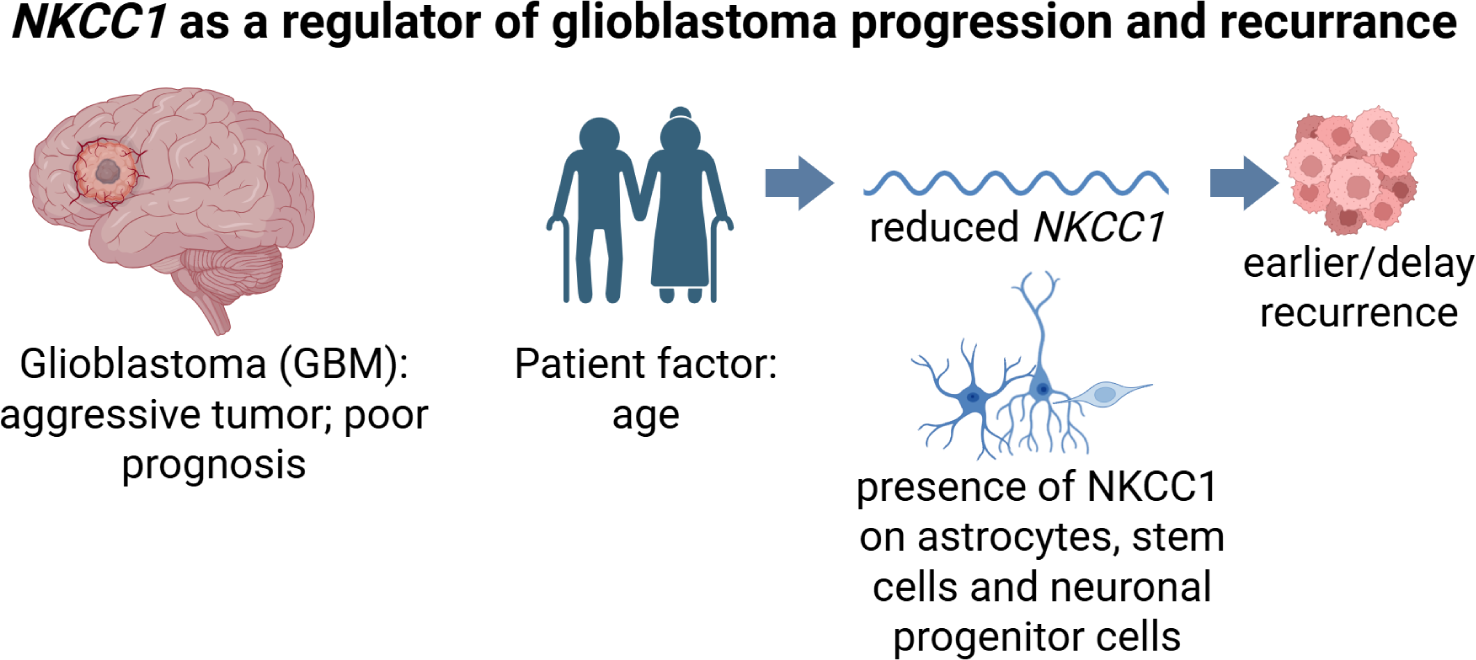

## Introduction

Glioblastoma (GBM) is the most common and most aggressive primary brain tumor in adults [1]. Despite intensive multimodal treatment approaches, including surgical resection, radiotherapy, and chemotherapy with temozolomide, the prognosis for patients remains extremely poor. The median survival time is approximately 15 months after diagnosis [2] and the five-year survival rate is less than 5% [3]. This unfavorable outcome is largely driven by the high invasiveness of the tumor, its genetic and epigenetic heterogeneity, and the ability of tumor cells to evade therapeutic strategies and develop resistance. Consequently, a central challenge in glioblastoma research is to identify molecular mechanisms that regulate tumor growth, invasiveness, and therapy resistance in order to develop novel therapeutic strategies. In this context, ion channels and transporters have received increasing attention, as they are indispensable for maintaining cellular homeostasis, regulating cell volume, and mediating signal transduction.

Among these, the cation-chloride cotransporters NKCC1 and KCC2 are of particular interest. They are essential regulators of intracellular chloride concentration and play a pivotal role in the functional development of the central nervous system. NKCC1 is mainly active in immature neurons, where it increases intracellular chloride levels, whereas KCC2 is expressed in mature neurons and lowers chloride concentration by extruding ions [4, 1, 5]. Through their opposing functions, these transporters shape the action of the neurotransmitter GABA, which depolarizes immature neurons but hyperpolarizes mature ones [6-9].

In GBMs, dysregulation of chloride transport appears to contribute directly to tumor progression. NKCC1, in particular, has emerged as a critical factor in glioma biology [10, 11, 5, 12-14]. By mediating chloride influx, NKCC1 facilitates changes in cell volume and cytoskeletal organization that are essential for GBM cells to migrate through the dense extracellular matrix of the brain. Upregulated NKCC1 activity has been associated with enhanced proliferation, invasiveness, and survival of GBM cells [4]. Pharmacological inhibition of NKCC1 has been shown in experimental models to reduce glioma cell migration and invasiveness, highlighting its potential clinical relevance [15-17]. Given its dual role in regulating both chloride homeostasis and glioblastoma cell behavior, NKCC1 represents a promising candidate for therapeutic intervention aimed at limiting tumor progression and improving patient outcomes.

The role of NKCC1 in GBM and its potential therapeutic implications highlight the relevance of these transporters for neuronal function and the pathogenesis of neurological and oncological diseases.

## Materials and Methods

### Human samples

The transcription analyses were performed with 61 glioblastoma samples surgically obtained from the Department of Neurosurgery at the University Hospital Jena. 10 FFPE samples were used for localization and co-localization of specific cells after immunofluorescence staining and confocal microscopy, which were provided by the Institute of Neuropathology, Charité-University Berlin. All tissue samples were histopathologically classified according to the current 2016 WHO classification for tumors of the central nervous system by the Institute of Neuropathology at Charitè Berlin, University Berlin [18]. All patients provided informed consent for the pseudonymized use of the extracted tumor material. The study was approved by the local ethics committee for human research (*Reg*.*-Nr. 2019-1400*).

### Total RNA extraction, reverse transcription and quantitative polymerase chain reaction

Surgically removed tissue was immediately stored at -80°C until processing. Total RNA from tissue sample was isolated, lysed in 1 ml Qiazol (Qiagen GmbH, Hilden, Germany) and homogenized with TissueLyser LT (Qiagen GmbH, Hilden, Germany) at 50 Hz for 5 min. The lysed tissue sample was mixed with 0.2 volume units (PU) of chloroform, centrifuged at 12,000 x g at 4 °C for 15 minutes. The upper aqueous RNA phase was removed and mixed with 1 PU isopropanol and centrifuged at 12,000 x g at 4 °C for 10 minutes. The resulting pellet was dissolved twice in 75 % ethanol and precipitated at -20 °C for one hour. The resulting pellet was air-dried for 10 minutes and resuspended in 40 µl RNASE-free water.

The RNA quality and concentration was determined spectrophotometrically (NanoDrop2000; Thermo Scientific, Wilmington, DE, USA). Complementary DNA synthesis was performed using the GoScriptTM Reverse Transcription System (Promega Corp., Mannheim, Germany) in a volume of 20 µl per reaction according to the manufacturer’s instructions.

The prepared cDNA was used as the basis for qPCR with the DyNAmo Flash SYBR Green qPCR Kit (Thermo Scientific Inc., Wilmington, DE, USA) to determine the mRNA levels of the transporters *NKCC1* and *KCC2*. The human-specific primers used for qPCR were designed based on mRNA coding sequences (NCBI Nucleotide) with NetPrimer (PREMIER Biosoft, Palo Alto, CA, USA) and NCBI Primer-BLAST and checked for hairpins and other secondary structures. The specificity of the amplification reaction was analyzed by gel electrophoresis and melting curve analysis.

The primer sets are listed in Table 1. The specific transcripts were amplified in a RotorGeneG (Qiagen GmbH, Hilden, Germany) with the following program: 7 min polymerase activation, 40 amplification cycles (95°C for 10s, 55°C for 20s, 72°C for 30s). To quantify the expression of target genes, we normalized the Ct value using *RPL13A* and *CYC1* as reference genes and normalized expression using the efficiency-based ΔCt method (Pfaffl, 2001).

**Table 1.**
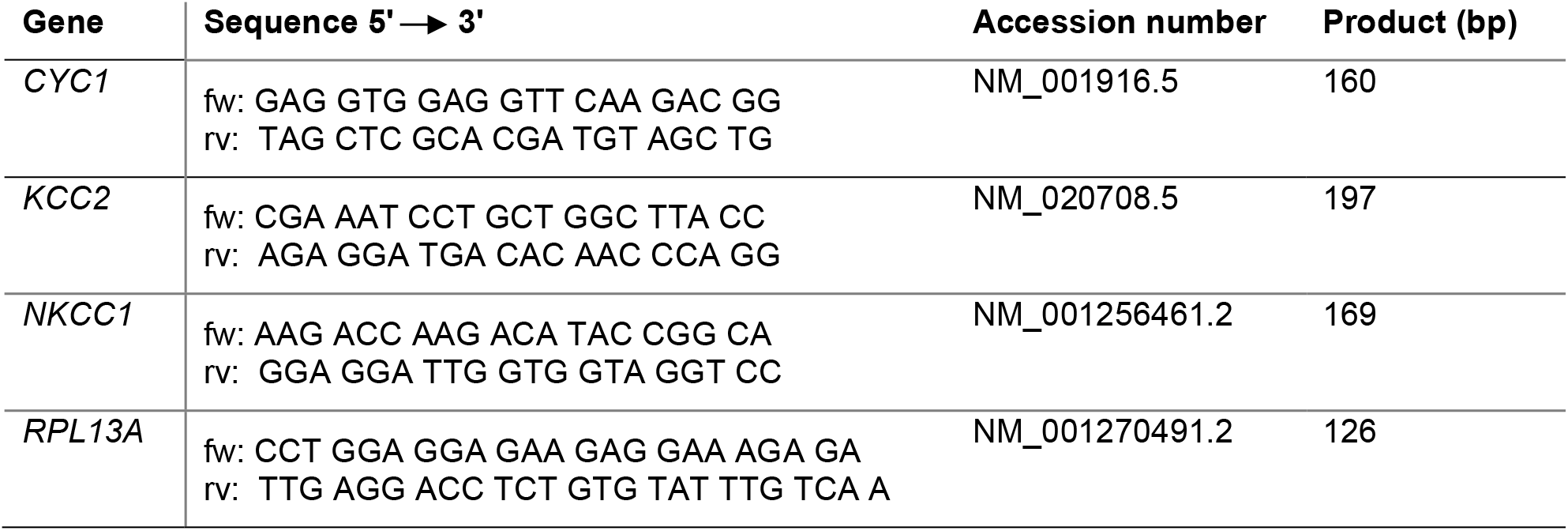
qPCR primers (fw: forward, rv: reverse)

### Immunofluorescence staining

The FFPE sections were deparaffinized by a descending ethanol series. Antigen retrieval was performed by incubating citrate buffer (pH 6.0) at 60 °C for 30 minutes. After washing with phosphate buffered saline (PBS), the non-specific binding sites were blocked with 20 % normal donkey serum (NDS) for 120 min. Hybridization of the primary antibodies: rabbit anti-NKCC1 (Proteintech™; 1:250), chicken anti-DCX (Abcam; 1:500), mouse anti-nestin (Abcam; 1:500), mouse anti-NeuN (Merck Millipore KGaA; 1:500) and/or mouse anti-S100b (Sigma Aldrich Chemie GmbH; 1:500) was performed overnight at 4 °C. The next day the slides were incubated at 4 °C. After two washes, the secondary antibodies were hybridized: Alexa488 donkey anti-mouse (Jackson Immunoresearch; 1:200), Alexa488 donkey anti-chicken (Jackson Immunoresearch; 1:200), Cy5 donkey anti-mouse (Jackson Immunoresearch; 1:200) and Rhodamine donkey anti-rabbit (Jackson Immunoresearch; 1:200) for 2 hours. Nuclear staining was performed with DAPI.

### Colocalization of NKCC1 with stem cell, glial, progenitor and neuronal marker using immunofluorescence staining

Ten GBM samples were used for fluorescent immunohistochemistry. Tissue sections were pre-embedded in paraffin and provided by Institute of Neuropathology, Charité-University Berlin. Colocalization analysis was performed for NKCC1 with markers of stem cells (Nestin), mature astrocytes (S100b), neuronal progenitor cells (Doublecortin, DCX), and mature neurons (NeuN).

Fluorescence imaging was performed using the Keyence microscope at 4× magnification for overview images. High-magnification images (40×) were acquired using the LSM980 confocal microscope (Zeiss, Jena, Germany). Individual cells were also imaged to confirm the specificity of the staining.

### Statistics

Statistical analyses were carried out using SPSS software (version 23). Gene expression (qPCR) data were first tested for normal distribution. Depending on the distribution, differences between two groups were analyzed using the Mann–Whitney U test.

For survival analysis, receiver operating characteristic (ROC) curves were generated in SPSS, and an optimal cutoff value was determined using the Youden index (J = sensitivity + specificity – 1). This cutoff was applied to stratify normalized expression values for subsequent analyses. Progression-free survival (PFS) and overall survival (OS) were evaluated using Kaplan–Meier survival analysis. A p-value of ≤ 0.05 was considered statistically significant.

## Results

### Age-dependent increase of NKCC1 in Glioblastoma

The patients’ age, sex, and MGMT methylation status were analyzed (Fig. 1). Significant differences were observed in *NKCC1* expression based on age: younger patients (≤ 60 years) exhibited approximately threefold lower NKCC1 expression (mean = 0.02374) compared to older patients (> 60 years) (mean = 0.07220), which was statistically significant (p = 0.027) (Fig. 1A/B).

**Fig. 1.**
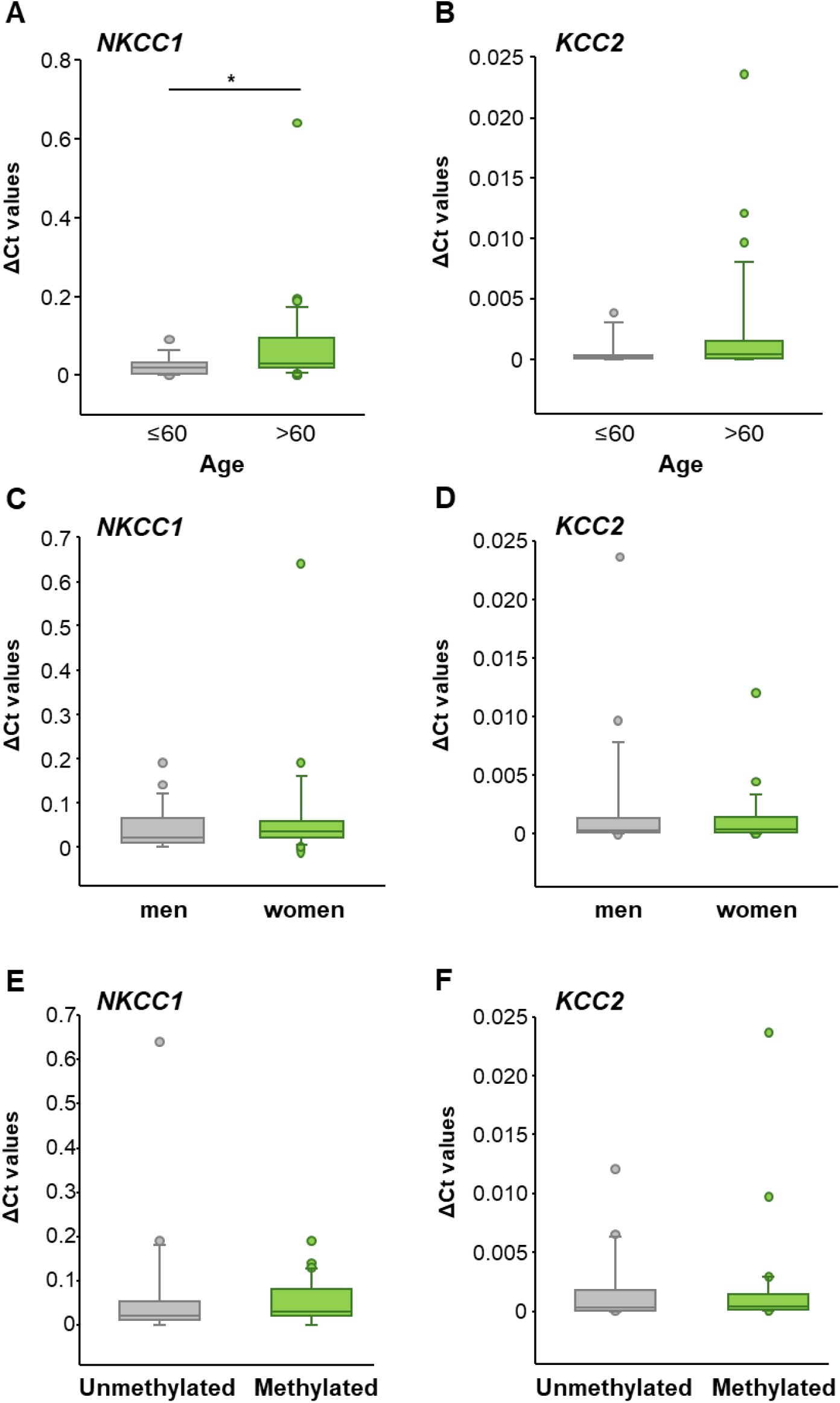
**A** *NKCC1* was expressed significantly more strongly in individuals over 60 years of age compared to those under 60. **B** There were no age-related differences in *KCC2* expression in glioblastoma The expression of *NKCC1* and *KCC2* was not affected by **C/D** sex or **E/F** methylation status. Mann–Whitney U test, \**p ≤* 0.05.

Regarding sex, *NKCC1* and *KCC2* expression levels were similar between males and females (*NKCC1*: males mean = 0.04089, females mean = 0.06978; *KCC2*: males mean = 0.00213, females mean = 0.00129) (Fig. 1C/D). Likewise, no significant differences were found based on MGMT methylation status (*NKCC1*: unmethylated promoter mean = 0.06367, methylated promoter mean = 0.04881; *KCC2*: unmethylated promoter mean = 0.00164, methylated promoter mean = 0.00175) (Fig. 1E/F).

### Decreased NKCC1 levels correlate with progression-free survival

To assess the impact of *NKCC1* expression on recurrence-free survival, a cutoff value of NE = 0.01401 was determined for individual patients (n = 48) using ROC analysis (Fig. 2A). Kaplan–Meier analysis demonstrated that patients with *NKCC1* expression below this cutoff (NE < 0.01401) had significantly longer recurrence-free survival during the first nine months after surgery compared to patients with higher *NKCC1* expression (NE > 0.01401; p = 0.019) (Fig. 2A).

**Fig. 2.**
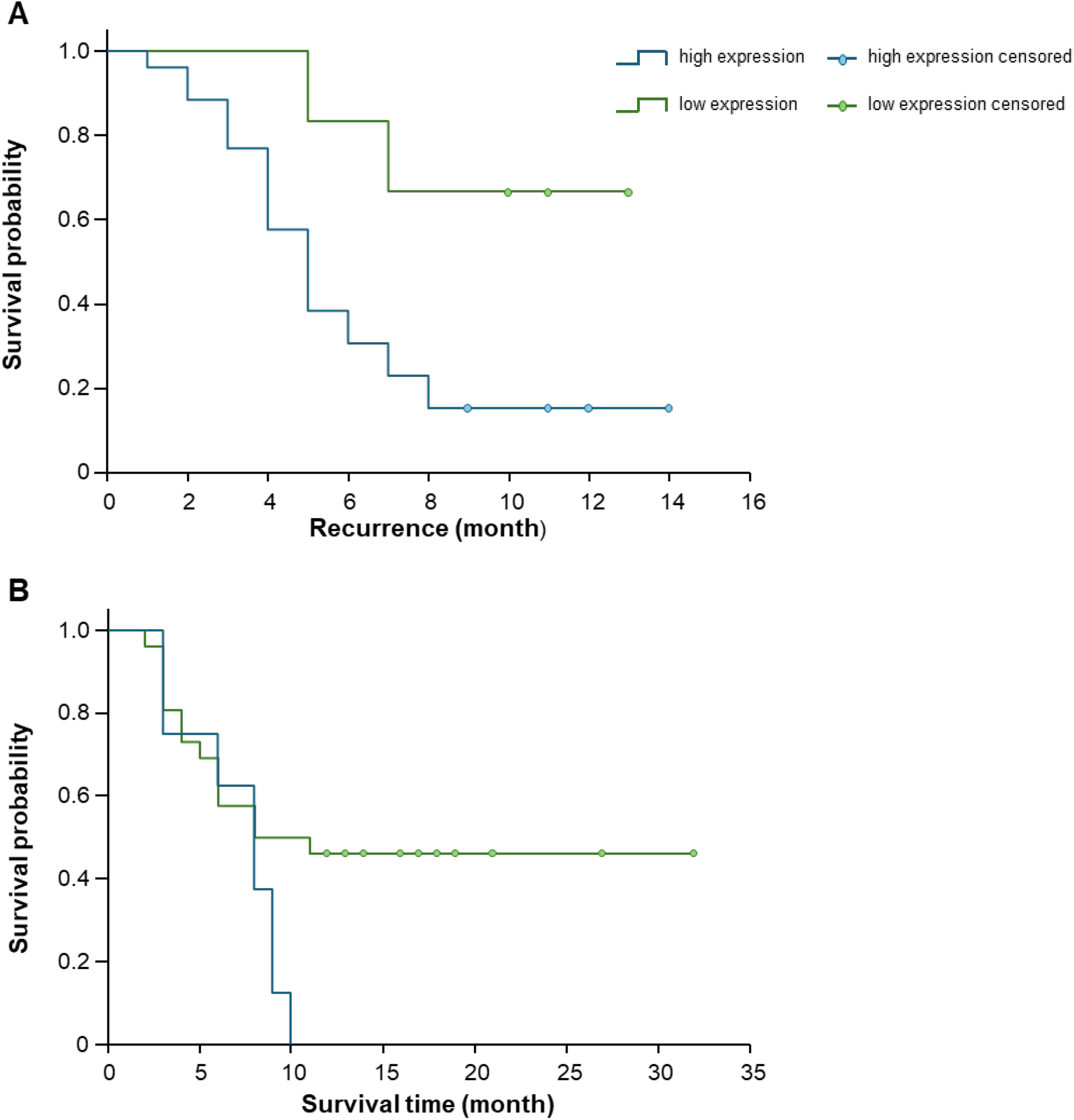
Progression-free and overall survival in relation to *NKCC1* expression. **A** Kaplan–Meier analysis showing that patients with low *NKCC1* expression had significantly longer progression-free survival after 9 months (*p* ≤ 0.05). **B** Overall survival tended to be longer in patients with low *NKCC1* expression after 12 months, but this difference was not statistically significant.

For overall survival analysis, 43 patients were included (16 patients excluded due to missing death data), and a cutoff value of NE = 0.04028 was established (Fig. 2B). Kaplan–Meier analysis revealed no significant association between *NKCC1* expression and overall survival (log-rank test; p = 0.078). Kaplan-Meier analysis based on gender revealed no differences in both overall survival and progression-free survival. In contrast, when stratified by age, the analysis showed differences in overall survival but not in progression-free survival.

### NKCC1 Colocalization with Specific Glial and Progenitor Cell Types in Glioblastoma

Ten GBM samples were subjected to fluorescence staining and subsequently imaged at both 4x and 40x magnifications for visual evaluation. NKCC1 was stained alongside various glial and neuronal markers, including Nestin, S100b, DCX, and NeuN. Nestin served as a marker for neural stem cells, S100b for astrocytes, DCX for neuronal progenitor cells, and NeuN for mature neurons.

Overview images at 4x magnification revealed a heterogeneous distribution of the labeled cell types throughout the tissue (Fig. 3). Rather than a uniform pattern, all samples displayed distinct regional clusters of specific cell populations. The fluorescence intensity of each marker protein (Nestin, S100b, DCX, or NeuN) was notably increased in defined areas, indicating local accumulations of the respective cell types.

High-resolution imaging at 40x magnification demonstrated colocalization of the chloride cotransporter NKCC1 with multiple cell types within the GBM tissue (Fig. 3). A pronounced overlap was observed between NKCC1 signals and Nestin-positive cells, suggesting expression in neural stem cells. Additionally, colocalization occurred with S100b-positive astrocytes and DCX-positive neuronal progenitor cells. Conversely, no significant colocalization was detected between NKCC1 and NeuN-positive mature neurons.

**Fig. 3.**
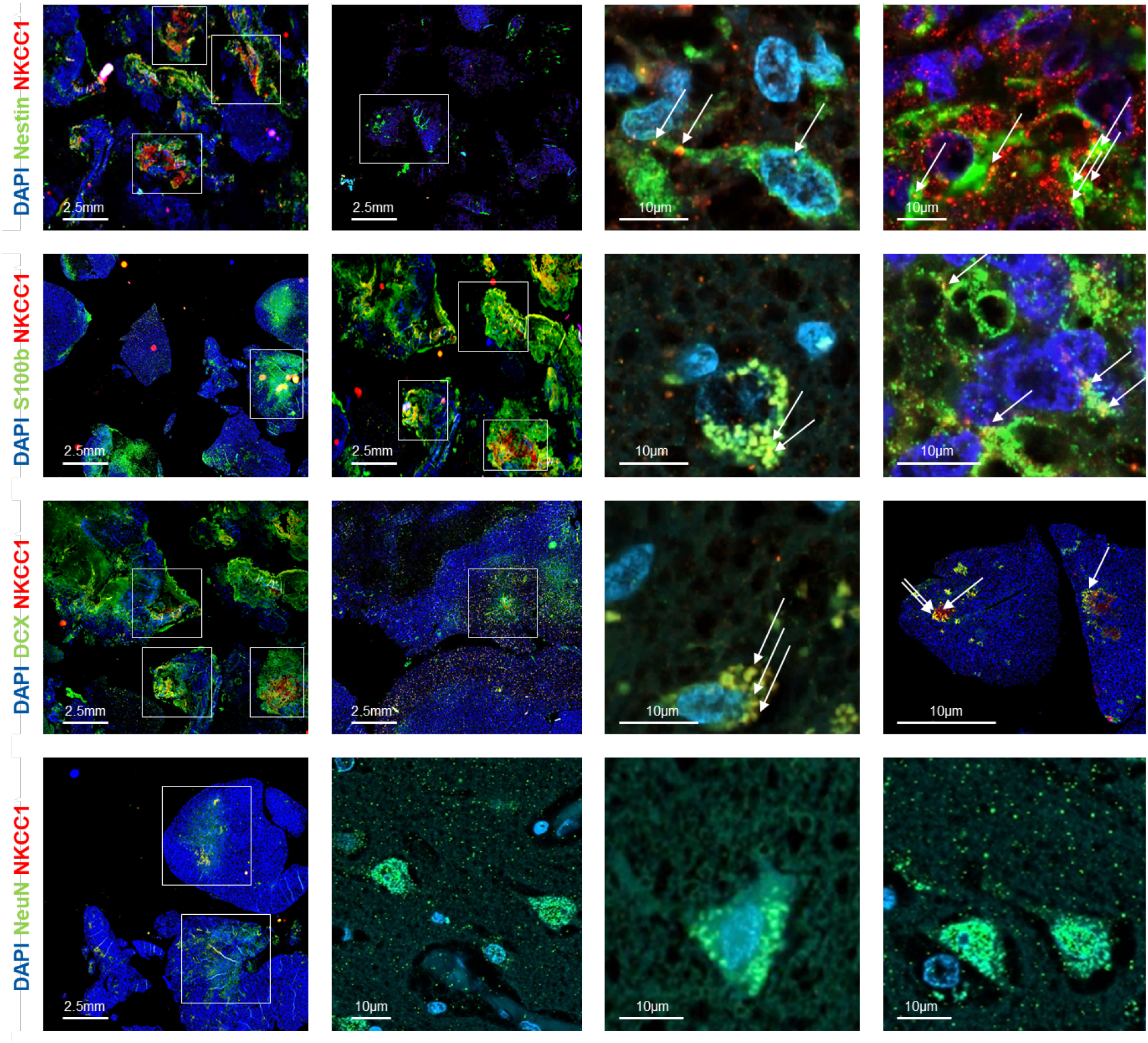
Immunohistochemical analysis of glioblastoma samples showing NKCC1 expression in different cell types. The first two columns show stained GBM samples at 4x magnification. These overview images illustrate the clustering behavior of different cell types: stem cells, astrocytes, neuronal progenitor cells, and mature neurons each co-stained with NKCC1. Clear colocalization of NKCC1 (red) was observed with Nestin-positive stem cells, S100b-positive astrocytes, and DCX-positive neuronal progenitor cells (green) and DAPI (blue). In contrast, no colocalization was detected with NeuN-positive mature neurons (green). Images in the third and fourth columns, acquired at 40x magnification, provide higher-resolution views of NKCC1 expression (red) in various cell types (green) within GBM tissue. White arrows mark the presence of NKCC1 in the different cell types.

## Discussion

The present study examined the expression and potential role of the chloride cation cotransporters NKCC1 and KCC2 in human glioblastomas, taking into account clinical parameters such as age, sex, and MGMT promoter methylation status. The results contribute to understanding the molecular heterogeneity of GBMs [19] and suggest possible implications for prognosis and therapy.

An age-dependent increase in *NKCC1* expression was observed, indicating a potential association between patient age and transporter activity. Elevated *NKCC1* may be linked to processes such as cell volume regulation and migratory capacity, which are relevant for tumor invasion [12]. This observation may partly explain the generally more aggressive tumor behavior observed in older patients [15, 5].

*NKCC1* also emerged as a potential marker for progression-free survival. Patients with lower *NKCC1* expression exhibited longer progression-free intervals, while overall survival was not significantly associated with *NKCC1* levels. These findings suggest that *NKCC1* gene expression may be particularly relevant for early tumor progression and recurrence rather than overall survival. From a translational perspective, *NKCC1* represents a potential therapeutic target. Current inhibitors, such as bumetanide, are limited by side effects and poor brain penetration, highlighting the need for the development of more selective and brain-penetrant compounds [15, 17].

Immunohistochemical analyses demonstrated cell type-specific NKCC1 expression in GBM tissue. NKCC1 colocalized with stem cell (Nestin), astrocyte (S100b), and neuronal progenitor cell (DCX) markers, but not with mature neurons (NeuN), consistent with its role in developmental and progenitor cell populations [20].

Overall, these results indicate a multifaceted role of NKCC1 in glioblastoma biology, particularly in tumor progression and recurrence. They support further investigation into targeting chloride homeostasis as a therapeutic strategy in GBM and provide a rationale for developing selective *NKCC1* inhibitors and evaluating the effects of modulating the NKCC1/KCC2 balance in vivo.

## Conclusions

This study offers new insights into the molecular heterogeneity of glioblastomas and identifies NKCC1 as a key regulator of tumor progression. The observed age-related increase of *NKCC1* underscores the importance of personalized treatment strategies that consider patient-specific molecular profiles. These findings suggest several promising avenues for future research, including the development of selective NKCC1 inhibitors with enhanced clinical applicability. In summary, this work advances our understanding of GBM biology and provides a foundation for developing more effective therapeutic approaches against this highly aggressive brain tumor.

## Author Contributions

All authors approved the final submitted version of the manuscript. A.T., D.F., M.H., S.K.: Data acquisition, data curation, and writing. C.S., F.S.: review & editing, final approval. S.K, D.F.: Conceptualization, writing, review, and editing.

## Acknowledgements

S.K. received support from Deutsche Forschungsgemeinschaft (DFG; grants KE1914/2-1; Interdisciplinary Center for Clinical Research Jena (Woman in science FF06); 2). A.T. was supported by the Interdisciplinary Center for Clinical Research Jena. Graphical abstract was created in BioRender. https://BioRender.com.

## References

1. Li K, Duan M, Lu Q, Liu J, He M & Zhang Y (2025) Advances in neuroscientific mechanisms and therapies for glioblastoma. iScience 28, 113347, doi: 10.1016/j.isci.2025.113347.

2. Stupp R, Hegi ME, Mason WP, van den Bent MJ, Taphoorn MJ, Janzer RC, Ludwin SK, Allgeier A, Fisher B, Belanger K, Hau P, Brandes AA, Gijtenbeek J, Marosi C, Vecht CJ, Mokhtari K, Wesseling P, Villa S, Eisenhauer E, Gorlia T, Weller M, Lacombe D, Cairncross JG, Mirimanoff RO, European Organisation for R, Treatment of Cancer Brain T, Radiation Oncology G & National Cancer Institute of Canada Clinical Trials G (2009) Effects of radiotherapy with concomitant and adjuvant temozolomide versus radiotherapy alone on survival in glioblastoma in a randomised phase III study: 5-year analysis of the EORTC-NCIC trial. Lancet Oncol 10, 459–466, doi: 10.1016/S1470-2045(09)70025-7.

3. Ostrom QT, Price M, Neff C, Cioffi G, Waite KA, Kruchko C & Barnholtz-Sloan JS (2022) CBTRUS Statistical Report: Primary Brain and Other Central Nervous System Tumors Diagnosed in the United States in 2015–2019. Neuro Oncol 24, v1–v95, doi: 10.1093/neuonc/noac202.

4. Cong D, Zhu W, Kuo JS, H. S & Sun D (2015) Ion transporters in brain tumors. Curr Med Chem 22, 1171–1181, doi: 10.2174/0929867322666150114151946.

5. Ma H, Li T, Tao Z, Hai L, Tong L, Yi L, Abeysekera IR, Liu P, Xie Y, Li J, Yuan F, Zhang C, Yang Y, Ming H, Yu S & Yang X (2019) NKCC1 promotes EMT-like process in GBM via RhoA and Rac1 signaling pathways. J Cell Physiol 234, 1630–1642, doi: 10.1002/jcp.27033.

6. Nascimento AA, Pereira-Figueiredo D, Borges-Martins VP, Kubrusly RC & Calaza KC (2024) GABAergic system and chloride cotransporters as potential therapeutic targets to mitigate cell death in ischemia. J Neurosci Res 102, e25355, doi: 10.1002/jnr.25355.

7. Peerboom C & Wierenga CJ (2021) The postnatal GABA shift: A developmental perspective. Neurosci Biobehav Rev 124, 179–192, doi: 10.1016/j.neubiorev.2021.01.024.

8. Pressey JC, de Saint-Rome M, Raveendran VA & Woodin MA (2023) Chloride transporters controlling neuronal excitability. Physiol Rev 103, 1095–1135, doi: 10.1152/physrev.00025.2021.

9. Schulte JT, Wierenga CJ & Bruining H (2018) Chloride transporters and GABA polarity in developmental, neurological and psychiatric conditions. Neurosci Biobehav Rev 90, 260–271, doi: 10.1016/j.neubiorev.2018.05.001.

10. Demian WL, Persaud A, Jiang C, Coyaud E, Liu S, Kapus A, Kafri R, Raught B & Rotin D (2019) The Ion Transporter NKCC1 Links Cell Volume to Cell Mass Regulation by Suppressing mTORC1. Cell Rep 27, 1886–1896 e1886, doi: 10.1016/j.celrep.2019.04.034.

11. Garzon-Muvdi T, Schiapparelli P, ap Rhys C, Guerrero-Cazares H, Smith C, Kim DH, Kone L, Farber H, Lee DY, An SS, Levchenko A & Quinones-Hinojosa A (2012) Regulation of brain tumor dispersal by NKCC1 through a novel role in focal adhesion regulation. PLoS Biol 10, e1001320, doi: 10.1371/journal.pbio.1001320.

12. Schiapparelli P, Guerrero-Cazares H, Magana-Maldonado R, Hamilla SM, Ganaha S, Goulin Lippi Fernandes E, Huang CH, Aranda-Espinoza H, Devreotes P & Quinones-Hinojosa A (2017) NKCC1 Regulates Migration Ability of Glioblastoma Cells by Modulation of Actin Dynamics and Interacting with Cofilin. EBioMedicine 21, 94–103, doi: 10.1016/j.ebiom.2017.06.020.

13. Sun H, Long S, Wu B, Liu J & Li G (2020) NKCC1 involvement in the epithelial-to-mesenchymal transition is a prognostic biomarker in gliomas. PeerJ 8, e8787, doi: 10.7717/peerj.8787.

14. Wang JF, Zhao K, Chen YY, Qiu Y, Zhu JH, Li BP, Wang Z & Chen JQ (2021) NKCC1 promotes proliferation, invasion and migration in human gastric cancer cells via activation of the MAPK-JNK/EMT signaling pathway. J Cancer 12, 253–263, doi: 10.7150/jca.49709.

15. Haas BR & Sontheimer H (2010) Inhibition of the Sodium-Potassium-Chloride Cotransporter Isoform-1 reduces glioma invasion. Cancer Res 70, 5597–5606, doi: 10.1158/0008-5472.CAN-09-4666.

16. Huberfeld G, Blauwblomme T & Miles R (2015) Hippocampus and epilepsy: Findings from human tissues. Rev Neurol (Paris) 171, 236–251, doi: 10.1016/j.neurol.2015.01.563.

17. Ilkhanizadeh S, Sabelstrom H, Miroshnikova YA, Frantz A, Zhu W, Idilli A, Lakins JN, Schmidt C, Quigley DA, Fenster T, Yuan E, Trzeciak JR, Saxena S, Lindberg OR, Mouw JK, Burdick JA, Magnitsky S, Berger MS, Phillips JJ, Arosio D, Sun D, Weaver VM, Weiss WA & Persson AI (2018) Antisecretory Factor-Mediated Inhibition of Cell Volume Dynamics Produces Antitumor Activity in Glioblastoma. Mol Cancer Res 16, 777–790, doi: 10.1158/1541-7786.MCR-17-0413.

18. Louis DN, Perry A, Wesseling P, Brat DJ, Cree IA, Figarella-Branger D, Hawkins C, Ng HK, Pfister SM, Reifenberger G, Soffietti R, von Deimling A & Ellison DW (2021) The 2021 WHO Classification of Tumors of the Central Nervous System: a summary. Neuro Oncol 23, 1231–1251, doi: 10.1093/neuonc/noab106.

19. Soeda A, Hara A, Kunisada T, Yoshimura S, Iwama T & Park DM (2015) The evidence of glioblastoma heterogeneity. Sci Rep 5, 7979, doi: 10.1038/srep07979.

20. Luo L, Guan X, Begum G, Ding D, Gayden J, Hasan MN, Fiesler VM, Dodelson J, Kohanbash G, Hu B, Amankulor NM, Jia W, Castro MG, Sun B & Sun D (2020) Blockade of Cell Volume Regulatory Protein NKCC1 Increases TMZ-Induced Glioma Apoptosis and Reduces Astrogliosis. Mol Cancer Ther 19, 1550–1561, doi: 10.1158/1535-7163.MCT-19-0910.

